# Dynamic Gating Mechanism of the b^0,+^AT-Mediated Arg Transport: Insights from ASMD Simulations

**DOI:** 10.64898/2026.06.03.729795

**Authors:** Qinjie Tang, Xing Zhang, Xuhua Li, Jing Dong, Haojia Li, Youshen Wu, Zhiwei Yang, Lei Li, Xinzhe Yu, Lei Zhang, Shengli Zhang

## Abstract

Heteromeric amino acid transporters (HATs) mediate essential amino acid flux across membranes, but the molecular dynamics of substrate translocation remain poorly defined for many family members. Here, using conventional and adaptive steered molecular dynamics (cMD and ASMD) simulations, we identify residue W230 in the b^0,+^AT transport channel as a dynamic gate that regulates arginine (Arg) influx through side chain flipping. By integrating dynamic network analysis with dynamical cross-correlation of residue motions, we show that regulatory signals propagate from the Arg binding site through transmembrane helix 5 (TM5), a connecting loop, and TM6 to reach W230. We propose a dynamic gating mechanism for b^0,+^AT - mediated amino acid transport. Arg binding at V186 triggers signal propagation that enhances cooperative interactions between W230 and Arg, driving the side chain flipping of W230. Our findings reveal a dynamic gating mechanism underlying b^0,+^AT - dependent Arg transport and suggest that residue-triggered side chain reorientation may represent a conserved and efficient strategy in transporter function.

**Author Summary:** Amino acids are the essential building blocks of life, and their transport across cell membranes is vital for nutrition and cellular signaling. Heteromeric amino acid transporters (HATs) mediate this process, yet how they physically move substrates through the protein at the atomic level remains poorly understood. In this study, we used advanced computer simulations to observe, in unprecedented detail, how b^0,+^AT—a key HAT member—transports the amino acid arginine. Our simulations revealed that a single residue, tryptophan 230 (W230), functions as a molecular gate: its side chain flips open to allow arginine to pass and then closes behind it, ensuring one-way traffic into the cell. We further discovered that the initial binding of arginine sends a signal through specific structural elements (helices and loops) to trigger this gate opening. This work not only uncovers a dynamic gating mechanism for b^0,+^AT but also suggests that similar side-chain flipping events may represent a common and efficient strategy used by other transporters to control substrate movement. Our findings provide a new framework for understanding transporter function and could inform future drug design targeting these critical membrane proteins.

## 1. Introduction

Moving arginine across a lipid bilayer is not trivial – it requires a finely tuned molecular gate. The heteromeric amino transporter b^0,+^AT performs this task with high fidelity, yet the dynamic events that control substrate passage have remained unclear. b^0,+^AT belongs to the HAT family, where a heavy chain and a light chain assemble into functional heterodimers[1–10]. These complexes primarily mediate the influx of cystine and cationic amino acids and the efflux of neutral amino acids in renal and intestinal epithelial cells[11].

In the b^0,+^AT-rBAT complex, loss-of-function mutations of either subunit disrupt transport activity, leading to cystinuria[12,13]. The inward-open conformation of the human b^0,+^AT-rBAT complex (PDB ID: 6LI9) exhibits a dimeric architecture stabilized by the rBAT subunit[14,15]. A previous study suggested that W230 forms a tight van der Waals contacts during substrate binding[6]. However, whether W230 primarily restricts the diffusion of small nonpolar amino acids across the membrane or also provides an additional electrostatic interaction site for substrate coordination has remained unclear and requires further investigation[6].

In this study, we investigated the dynamic process of Arg transmembrane transport mediated by b^0,+^AT. We first performed extensive long-timescale conventional molecular dynamic (cMD) simulations (3×1000 ns) to establish a stable conformational ensemble of the b^0,+^AT-Arg complex. Because amino acid transport across membranes is often driven by proton or ion gradients[5,10], which provide an effective pulling force, we subsequently employed adaptive steered molecular dynamics (ASMD) simulations to drive Arg translocation through the complex embedded in a POPC membrane.

By integrating potential of mean force (PMF) analysis, force–displacement curves, binding free energy calculations, interaction fingerprints, dynamic network analysis, and mutational studies, we demonstrate that arginine transport is governed by a “dynamic gating mechanism”, with W230 serving as a key gating residue within the b^0,+^AT transport channel. Dynamic network analysis further traced the top 100 signaling pathways from Arg to W230, all originating from the direct interaction of Arg with residue V186. Our results reveal that the signal propagates through TM5, across a connecting loop, and into TM6, ultimately reaching W230. These dynamic network paths, together with the observed correlation between TM5 and TM6, suggest that arginine binding at V186 induces signal propagation that enhances mutual interactions between W230 and Arg, ultimately driving the side-chain flipping of W230. Such signal transmission and conformational rearrangement collectively underpin the “dynamic gating mechanism” of b^0,+^AT for arginine transport.

Beyond characterizing Arg transport via b^0,+^AT, our findings suggest a gating strategy that may be generalizable to other secondary active transporters. For instance, the conserved aromatic residue F252 in human LAT1 has been shown to form an extracellular occlusion gate[4,16]. The molecular dynamic gating mechanism of W230 offers a novel perspective on transporter gating, distinct from the well-characterized large-scale helical movements. While the latter involve major conformational rearrangements of transmembrane helices, side-chain flipping represents a localized and efficient regulatory strategy for controlling substrate passage.

## 2. Results

### 2.1. ASMD simulation and the characterization of the Arg transport channel

To investigate the dynamic process of Arg transmembrane transport mediated by b^0,+^AT, Three independent 1000-ns conventional MD simulations were performed to assess the structural stability of the b^0,+^AT–Arg complex (Figure S1, S2). The resulting stable conformational ensemble provides a reliable set of initial structures for subsequent ASMD simulations.

Based on the stable b^0,+^AT-Arg complex structure, we characterized the transport pathway of the Arg ligand using the ASMD simulation, an enhanced sampling simulation method.

As shown in Figure 1a, the PMF varies as a function of the pulling distance. The estimated energy barrier for moving the ligand from the binding pocket toward the extracellular side is approximately 140 kcal/mol. In contrast, the barrier toward the intracellular side is only 40 kcal/mol, which aligns with the inward-open conformation of the b^0,+^AT. While the absolute values of these computed barriers (40–140 kcal/mol) are significantly higher than the estimated free energy available from a proton-motive force per transport cycle (approximately 5 to 15 kcal/mol, depending on coupling stoichiometry)[5,10]. This discrepancy is expected for non-equilibrium steered simulations where high pulling rates introduce substantial dissipative work. Nevertheless, the pronounced asymmetry in the energy profile robustly indicates a directionally biased transport pathway, thereby supporting the validity of the structural model and the simulation approach.

**Figure 1.**
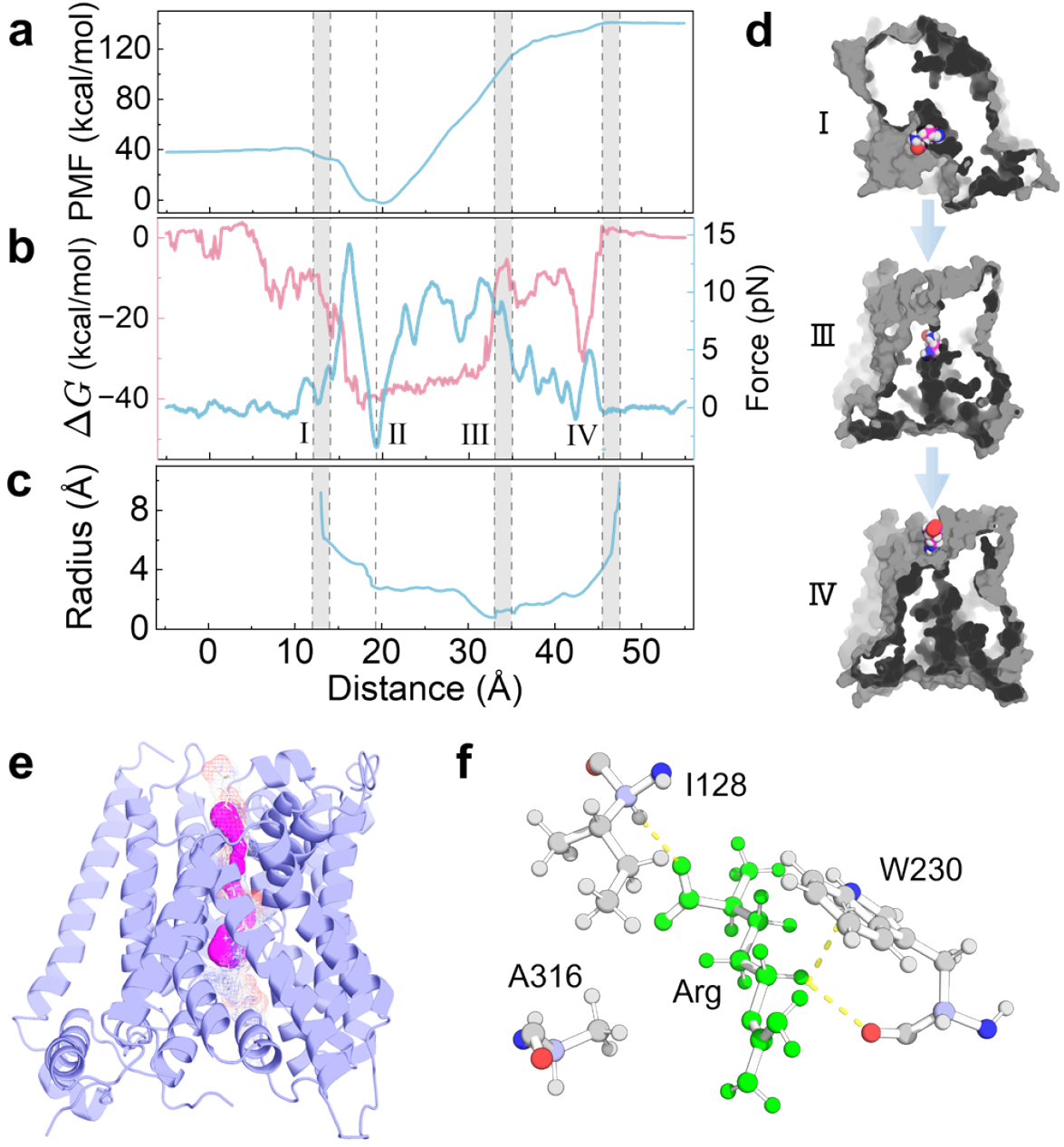
Process of Arg transport and analysis. (a) Variation of PMF with the reaction coordinate. The distance values increase from the intracellular side to extracellular side. (b) Comparative analysis of binding free energy *ΔG* and pulling force, where the pink line represents *ΔG* and the blue line represents pulling force. Region `I` denotes the position where the channel exits to the intracellular side, Region `II` is the ligand-binding pocket, Region `III` is the narrowest point of the transport channel, and Region `IV` is the channel entrance on the extracellular side. (c) Variation of the channel radius with the distance. In these pictures, regions `I`, `III`, and `IV` are marked in gray. (d) Three structural snapshots correspond to regions `I`, `III`, and `IV`. (e) Comparison of the channel for Arg movement revealed by ASMD simulation and the channel predicted by Caver 3.0. (f) Analysis of protein-ligand structural interactions at the narrowest point of the ligand transport channel, with Arg shown in green.

In physiological conditions, substrate translocation is driven by proton or ion gradients[5,10]. To mimic this driving force in our simulations, we applied an adaptive external pulling force in the ASMD simulations. The ASMD pulling force profile (blue line in Figure 1b) demonstrates that the maximum force occurs when the ligand transitions from the binding pocket toward the extracellular entrance. This suggests that the resistance is governed by both protein-ligand binding interactions and steric constraints from the narrow pore geometry.

The high PMF for extracellular exit (Figure 1a) stems mainly from overcoming steric hindrance in the pore; although, binding free energy also contributes substantially across the transport process. To evaluate this contribution, we computed the binding free energy along the pulling trajectory—sampling every 100 frames—and compared its evolution with the pulling force (Figure 1b). This integrated analysis identified four critical sites along the pathway:

- **Site** `IV`: Marked by the initial rise in both binding free energy and pulling force, indicating ligand entry from the extracellular side.
- **Site** `III`: Features minimal binding free energy (also see the reduced main interactions Fig 3a) but a peak in pulling force, suggesting dominance of steric resistance from the narrow pore.
- **Site** `II`: Corresponds to the ligand-binding pocket, with maximum binding free energy and high pulling force.
- **Site** `I`: Characterized by the decline of pulling force to zero and the lowest binding free energy, reflecting ligand release into the intracellular side.

These sites are consistent with the channel radius obtained from the HOLE2 analysis (Figure 1c). Structure snapshots of b^0,+^AT-Arg at Sites `I`, `III`, and `IV` are displayed in Figure 1d. The transport pathway derived from the ASMD trajectory is shown in Figure 1e, and is in strong agreement with the channel predicted by Caver3.0 software[17,18], further validating the accuracy of the identified pathway.

Given that Arg encounters significant steric hindrance at the narrowest point of the channel, clarifying the interactions at this location is crucial. As represented in Figure 1f, residues W230, I128, and A316 exhibit strong interactions with Arg, underscoring their importance in Arg transport.

### 2.2 Key residues for Arg transport identified by interaction analysis

As summarized in Table S1 (based on MM/PBSA calculations), W230 exhibited the strongest contribution to the binding free energy. To further investigate the functional role of residue W230, we analyzed its conformational changes during Arg transport (Figure 2a). The conformational evolution of W230 can be divided into three distinct stages as Arg traverses the channel: Initially, the indole ring of W230 laterally occupies and occludes the channel (Stage `I`). As Arg enters the constricted region, a pronounced flip of the W230 side chain opens the channel (Stage `II`). After the ligand moves into the binding pocket, the side chain reverts to its original position, thereby resealing the channel (Stage `III`). This “close–open–close” cycle functions as a dynamic gate that regulates Arg passage. Moreover, by promptly closing behind the translocating ligand, this gating mechanism ensures unidirectional transport toward the intracellular side, representing a proposed dynamic gating mechanism for Arg translocation.

**Figure 2.**
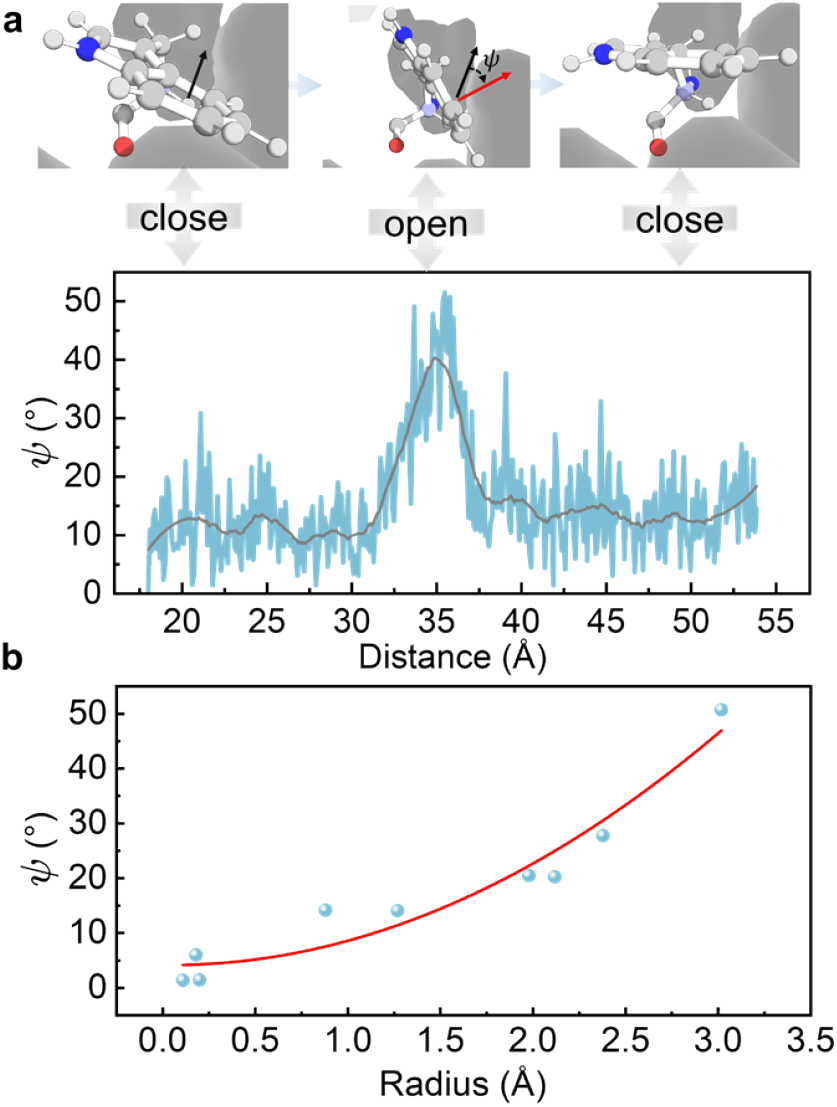
Analysis of the W230 side-chain flipping angle ψ. (a) Evolution of *ψ* as a function of the pulling coordinate (reaction coordinate). The close, open, and close states of the W230 side chain are schematically illustrated above. (b) Relationship between the angle *ψ* and the channel radius at Site `III`. Blue points represent data from the simulation trajectory, and the red solid line is the corresponding fitting curve.

To quantitatively characterize the side-chain flipping of W230, we defined an angle *ψ* based on the normal vector of the benzene ring plane relative to its initial position (Figure 2a). The variation in *ψ* showed an initial increase followed by a decrease, with a maximum reorientation of approximately 50°, which aligns well with the proposed gating mechanism during Arg transport (Figure 2a). Furthermore, we characterized the geometric impact of this flipping motion. Figure 2b presents the relationship between the angle (*ψ*) and the channel radius at Site `III`. The scatter plot demonstrates that the radius increases monotonically with *ψ*. We can see that the side-chain flipping of W230 expands the channel radius at the thinnest position Site `III`, allowing Arg to pass through the channel in b^0,+^AT.

### 2.3. Interaction fingerprint analysis of ASMD simulations

Our ASMD simulations revealed side-chain flipping of W230, suggesting its potential key role in Arg transport. To further explore the molecular mechanism underlying this flipping, we performed a systematic interaction fingerprint analysis of the ASMD trajectory[19]. We identified the top five key residues with the highest interaction frequencies and generated time-evolution maps for the hydrophobic, Hydrogen bonds, van der Waals, cation-π interactions (Figure 3a). A 2D interaction diagram further illustrates the specific interactions between these residues and the ligand (Figure 3b).

**Figure 3.**
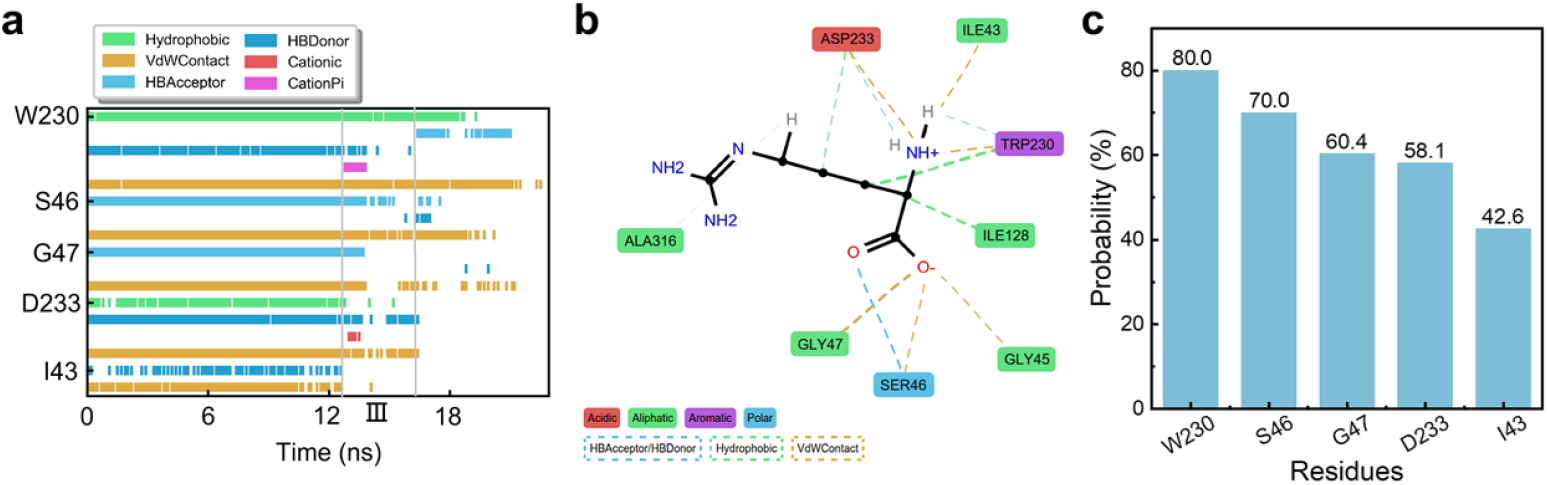
Interaction fingerprint analysis of the process of Arg transport through the narrowest region of the channel. (a) The time evolution of interactions for the five most frequently interacting residues. (b) Ligand-protein 2D interaction map. Key residues are colored according to physicochemical properties, interaction types are represented by the color of dashed lines, and the width of the line reflects the interaction duration. (c) The top five proportions of interaction time between the key residues and Arg.

Among the top five key residues, W230 maintained stable interactions with the ligand over 80% of the simulation time—more than any other residue—and exhibited diverse interaction modes. Other frequently interacting residues, such as Ser46, Gly47, and Asp233, displayed overall interaction frequencies above 50%, predominantly mediated by van der Waals contacts (Figure 3c).

Notably, a change of W230 from a hydrogen-bond donor to an acceptor was observed around 16 ns (site `III`), coinciding with the conformational flip of its side chain. Specifically, the NE atom of Arg likely forms a hydrogen bond with the backbone carbonyl oxygen of W230 (donor) initially. After the flip, the side-chain indole NH of W230 reorients to donate a hydrogen bond to one of Arg’s carboxylate oxygen atoms (acceptor). This switch in interaction mode may be triggered by a transient cation–π interaction occurring at ∼12 ns. We propose that as Arg approaches, its guanidinium group engages in a brief cation–π interaction with the electron-rich face of the W230 indole ring. This attractive interaction likely pulls Arg closer and perturbs the local electrostatic environment, promoting the subsequent side-chain rotation and hydrogen-bond role reversal. After the flip, the geometry changes, and the cation–π interaction dissipates. The high interaction frequency and multimodal profile of W230 support its role as a dynamic structural gate during Arg transport, facilitated by synergistic hydrophobic, van der Waals, and polar interactions.

These findings indicate that b^0,+^AT–Arg relies on a highly coordinated and diverse interaction network during substrate transport, with W230 serving as a key nodal residue enabling precise regulation through its dynamic interaction landscape.

### 2.4. Dynamic network path analysis

Using Bio3D, we computed the dynamical cross-correlation matrix (DCCM) and conducted dynamic network path analysis to investigate internal correlations and allosteric signal transduction during Arg translocation and W230 flipping[20–27] (Figure 4a,b). These signaling pathways are triggered by the direct interaction of Arg with residue V186 on TM5. Through dynamic network analysis, we identified the top 100 signaling pathways from Arg to W230, where the signal propagates through TM5, across a connecting loop, and into TM6 before finally reaching W230 (Figure 4b). The correlation values of key nodes a re provided in Table S2. Additionally, the DCCM indicates a strong negative correlation between TM5 and TM6 (Figure 4a), which typically reflects anti-correlated motions that may facilitate a gating-like opening/closing between these helices. By integrating dynamic network analysis with the observed negative correlation between TM5 and TM6, we propose that ligand binding at V186 triggers the signal propagation. This promotes cooperative interactions between W230 and Arg, leading to the flipping of W230.

**Figure 4.**
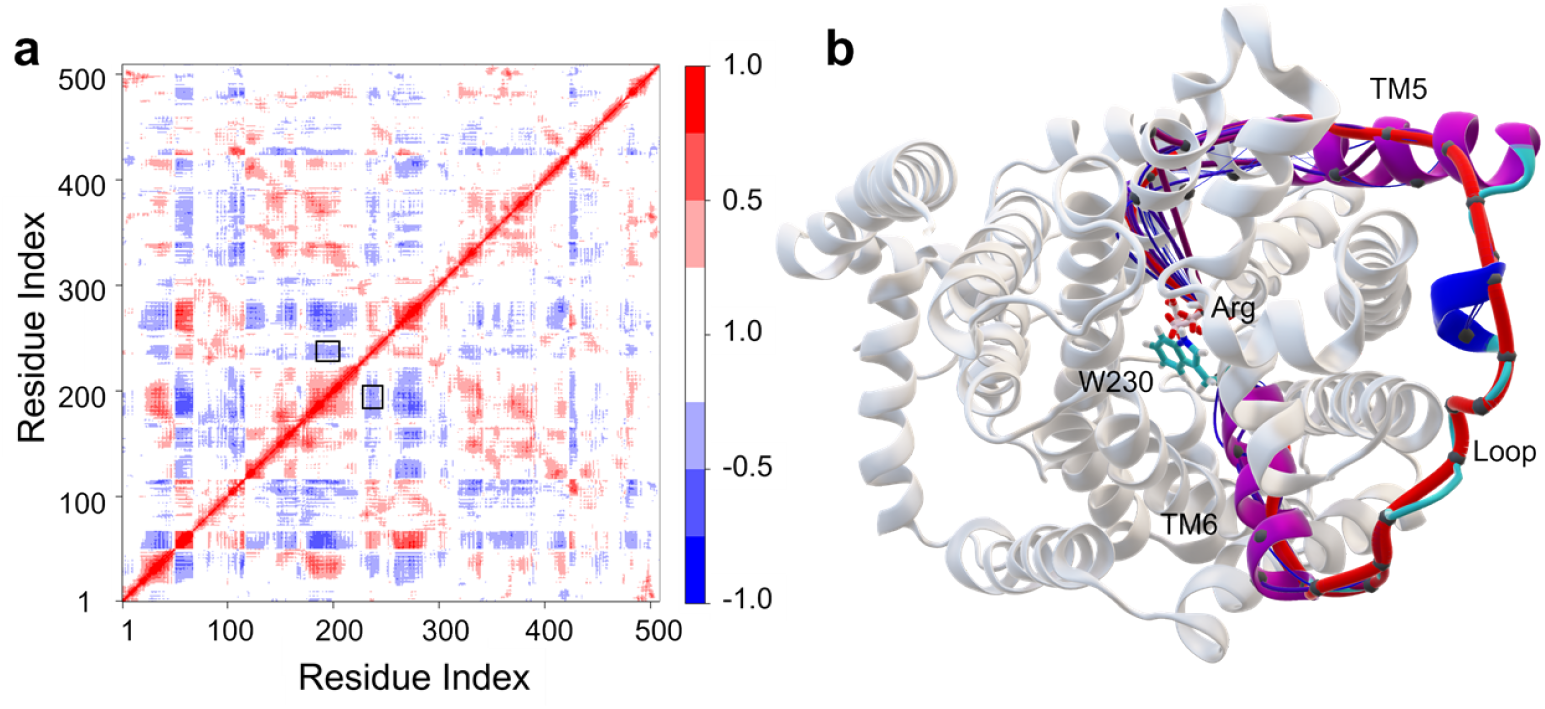
Correlation and the dynamic network path analysis. (a) DCCM analysis. The region of correlation between TM5 and TM6 is enclosed in a black rectangle. (b) Analysis of the signaling pathway from Arg to W230, with the optimal path highlighted by a thick red line, suboptimal paths by thin blue lines, and the constituent α-helix and loop regions colored purple and cyan, respectively.

### 2.5. Verification of key sites with W230A simulation

To validate the dynamic gating role of W230 in Arg transport, we introduced a W230A mutation and carried out comparative ASMD simulations to evaluate changes in the PMF and pulling forces. As shown in Figure 5a, the PMF values of the W230A mutant are lower (more favorable) than those of the wild type at nearly all positions, indicating a significantly reduced free energy barrier for Arg translocation in the mutant. Consistently, Figure 5b reveals a pronounced decrease in pulling force throughout the transport process, especially from the channel entrance to the ligand-binding pocket (corresponding to positions 20 Å– 45 Å). At labeled position III, the site of the W230 mutation, both PMF and force values were considerably lower than those in the wild type. These findings suggest that the W230A mutation not only alters PMF and force locally but also significantly affects the entire transmembrane transport process of Arg.

**Figure 5.**
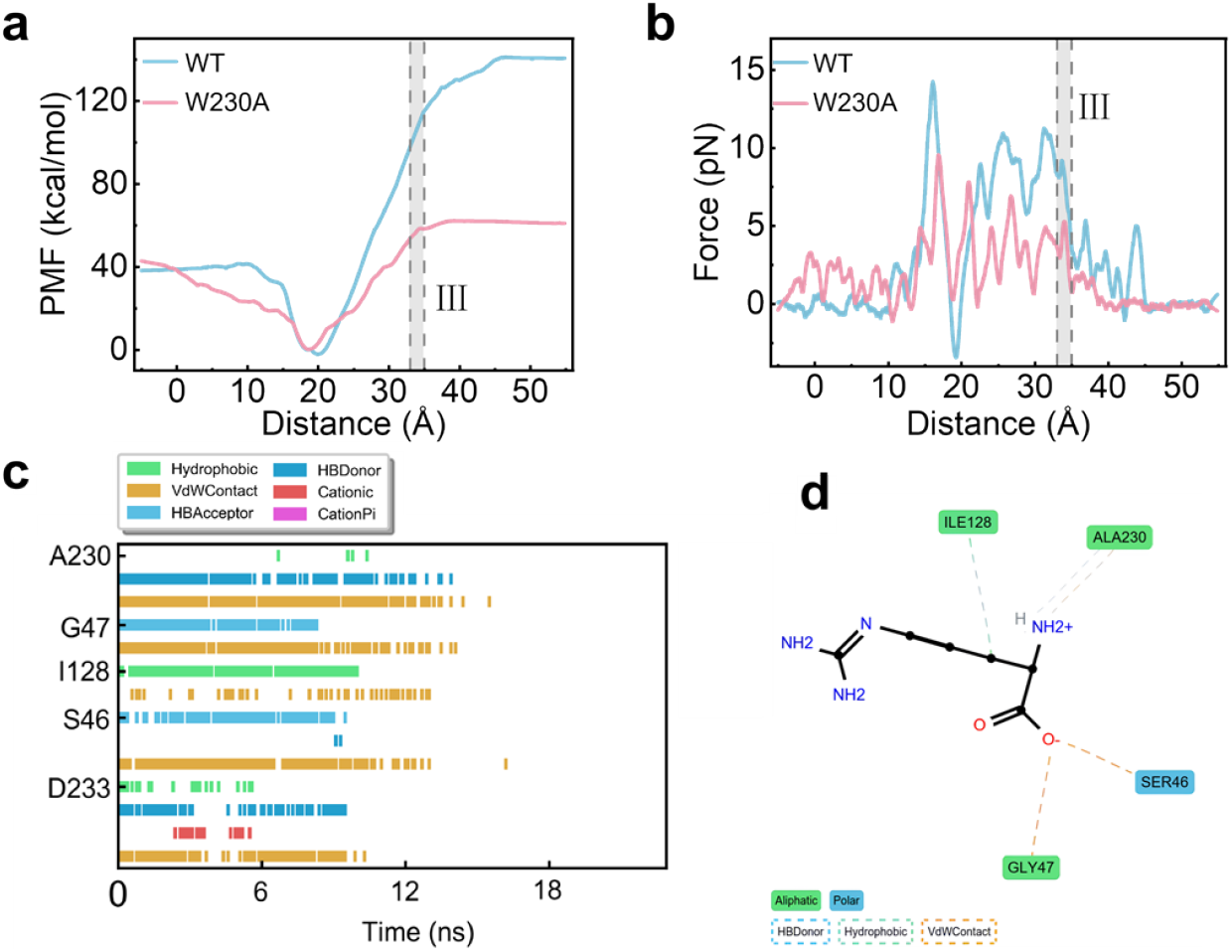
The PMF and pulling force comparation of W230A mutant with those of the wild-type and the interaction fingerprint analysis of W230A mutant. (a, b) Comparison of PMF and pulling force between WT and W230A. The blue line represents the WT, and the pink line represents the W230A. (c, d) Interaction fingerprint analysis of the Arg transport in the W230A mutant.

We further performed interaction fingerprint analysis on the W230A mutant (Figure 5c, d). Compared with the wild type (Figure 3), the mutant exhibited markedly reduced probabilities for multiple interaction types—including hydrophobic, van der Waals, and hydrogen-bond interactions (Figure 5c, d and Figure S5). In agreement with these computational results, experimental studies have reported that both the disease-related W230R mutation and the alanine-substitution W230A nearly abolish b^0,+^AT transport activity[14], providing direct support for our findings. Together, these data demonstrate that W230 is essential for facilitating ligand transport in the b^0,+^AT-Arg complex and corroborate its role in a dynamic gating mechanism.

This dynamic gating mechanism, particularly the side-chain flipping event, is consistent with previous reports[28–30]. For example, multiple transporter families employ analogous gating strategies to regulate substrate translocation across membranes. Such mechanisms are generally described by the “alternating access” model, in which transporters switch between inward-open and outward-open conformations to enable substrate passage, as documented for Major Facilitator Superfamily (MFS) transporters[30], glutamate transporters[28], and sodium-proton exchangers[29]. In our study, the transport process mediated by the heterologous amino acid transporter b^0,+^AT–Arg is strongly influenced by the side-chain flipping of W230, suggesting that it may also operate under a similar alternating-access mechanism. However, the detailed dynamic characteristics governing the transition between inward-open and outward-open states in b^0,+^AT remain to be fully elucidated.

## 3. Conclusion

In summary, this work systematically elucidates the transmembrane transport mechanism of arginine mediated by the b^0,+^AT–Arg complex using conventional and adaptive steered molecular dynamics simulations. We identified a well-defined transport channel within b^0,+^AT and characterized its translocation pathway through potential of mean force analysis, revealing W230 as a central gating residue. Integrating dynamic network analysis with cross-correlation of residue motions shows that arginine binding at V186 on TM5 triggers allosteric signal propagation through a connecting loop to TM6, ultimately inducing side-chain flipping of W230. This “close-open-close” cycle of W230 acts as a dynamic gate that not only permits arginine passage but also likely contributes to unidirectional transport by resealing the channel behind the ligand. The essential role of W230 was validated by mutational simulations (W230A), which severely impaired transport. Beyond b^0,+^AT, this dynamic gating strategy – based on localized aromatic side-chain flipping rather than large-scale helical rearrangements – may be conserved among other heteromeric amino acid transporters, such as LAT1 (F252). Our findings thus advance the mechanistic understanding of b^0,+^AT and propose ligand-triggered side-chain reorientation as an efficient and potentially generalizable regulatory principle in transporter biology.

## 4. Methods

### 4.1 Conventional Molecular Dynamics (cMD) Simulations

The simulation system was constructed based on the crystal structure of the human b^0,+^AT -rBAT complex bound to arginine (PDB ID: 6LI9)[14]. To focus on the transmembrane transport region, only the transmembrane domains (TMs) of b^0,+^AT and rBAT were retained; the extracellular domain of rBAT truncated to reduce system complexity. The protein-arginine complex was embedded into a periodic box using the CHARMM-GUI[31–38], associated with a 1-palmitoyl-2-oleoyl-sn-glycero-3-phosphocholine (POPC) lipid bilayer[14]. The system was hydrated with 150 mM NaCl solution layers (22.5 Å thick on each side), resulting in final box dimensions of 81 Å × 81 Å × 104 Å and 79826 atoms (Figure S1a).

In the initial structure, proteins used the AMBER FF19SB as the force field, and lipids used the Lipid21 force field. The solvent uses the OPC explicit water model[39–42].

The simulation protocol included three stages: energy minimization, NPT equilibration, and production cMD. Energy minimization was performed using the conjugate gradient method for 5000 steps to remove steric clashes. Six rounds of NPT equilibration (125,000 steps each, 1 fs time step) were conducted at 310 K to stabilize the lipid bilayer and solvent. In the dynamic simulation, temperature was controlled using the Langevin dynamics algorithm, and pressure was regulated via the Berendsen barostat method, with an integration time step of 2 fs. Three independent 1000 ns cMD simulations were then performed using the PMEMD[43–45] with cuda11.8. Trajectory analysis was performed using CPPTRAJ[46].

### 4.2 Adaptive Steered Molecular Dynamics (ASMD) Simulations

ASMD simulations were used to study the transmembrane transport of arginine[47–53]. Steered MD (SMD) applies a guiding force along a reaction coordinate to accelerate ligand dissociation or binding. The Jarzynski equality relates nonequilibrium work to free energy differences:

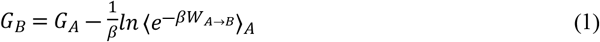

Where *G*_*B*_ and *G*_*A*_ are the free energy of state A and B, *β* = 1 *k*_*B*_*T, W*_*A*→*B*_ is the non-equilibrium work performed on the system during the ASMD simulation.

In SMD, many simulations must be run in order to converge the PMF. To improve convergence, ASMD divides the reaction path into stages. The equilibrated conformation was used as the initial structure, with the translocation distance of arginine from extracellular to intracellular as the reaction coordinate. The process was divided into 15 stages (4 Å each), with 2,000,000 steps per stage (2 fs time step). A pulling velocity of 1 Å/ns and a harmonic force constant of 5 kcal/mol·Å^2^ were applied (Figure S3, S4). Twenty-five independent trajectories were run per stage[54]. PMF was computed by combining all stages to obtain the free energy profile.

### 4.3 MM/GBSA Binding Free Energy Calculation

The molecular mechanics generalized Born surface area (MM/GBSA) method was used to calculate the binding free energies between b^0,+^AT and arginine during ASMD [54]. MM/GBSA is a widely recognized method and tool used for the assessment of free energy[55].

The binding free energies (Δ*G*_*bind*_) for the protein-ligand interactions were evaluated using the MM/GBSA approach implemented in AmberTools20[40,56], which has been proven effective in predicting binding affinities for a wide range of protein-ligand interactions[57,58]. Details of parameters are like those performed the previous works[59,60]. In brief, Δ*G*_*bind*_ was calculated using the equation:

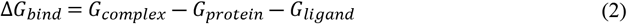

In addition to the overall binding free energy, the MM-GBSA decomposition method was used to calculate the binding interaction between the ligand and each residue in the protein. The contribution of each residue to the binding energy includes three terms: van der Waals contribution (Δ*E*_*vdw*_), electrostatic contribution (Δ*E*_*ele*_) in a vacuum, and solvation contribution (Δ*E*_*solvation*_):

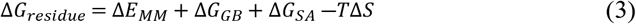

The molecular mechanical contribution Δ*E*_*MM*_ consists of electrostatic (Δ*E*_*ele*_) and van der Waals (Δ*E*_*vdw*_). Δ*G*_*solvation*_ includes denote polar energy Δ*G*_*GB*_ and nonpolar solvation free energy Δ*G*_*SA*_. −*T*Δ*S* represents the entropic contribution. All energy components were computed over 100 snapshots evenly extracted from each MD trajectory.

### 4.4 Conformational Analysis

Free energy landscape (FEL) analysis can provide information on the free energy basin, free energy barrier, and representative conformation of MD simulations. Representative conformations were extracted from low-energy regions and compared with the initial structure[14]. First, one needs to know the probability density *P(x)* of each conformation *{x}* and use the Boltzmann relation, free energy *G(x)* can be calculated:

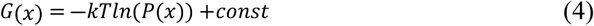

In this study the FEL was constructed using root mean square deviation (RMSD) of the protein backbone and radius of gyration (Rg)[61]. The GROMACS gmx sham command was used during the FEL analysis[62]. Structural plotting and visualization were accomplished by PyMOL.

### 4.5 Interacting Fingerprint Analysis

Interaction fingerprint (IFP) analysis was performed using ProLIF (Protein-Ligand Interaction Fingerprints) to probe the protein-arginine interactions throughout the ASMD simulations[63]. ProLIF digitally encodes molecular interactions into binary fingerprints and can process data from MD trajectories, crystal structures, or docking poses for various complex types. It detects interactions, including hydrogen bonds and hydrophobic contacts, using customizable geometric parameters defined by SMARTS patterns. The results were output in a pandas Data Frame, enabling statistical analysis and visualization (e.g., LigNetwork plots) to identify key residues.

### 4.6 Dynamic Network Path Analysis

To identify the allosteric pathway from Arg to W230, we employed dynamic network analysis as implemented in the Bio3D package[20–27]. In this approach, the dynamic cross-correlation matrix (DCCM) was converted into an adjacency matrix to construct a network where each node represents a Cα atom. The probability of information transfer along an edge was quantified by the weight, defined as −*log*|⟨*C*_*ij*_⟩|. A correlation threshold (*C*_*thr*_) was applied to filter out negligible fluctuations; only residue pairs with *C*_*ij*_ ≥ *C*_*thr*_ were connected. This method effectively reveals long-range communication pathways within a protein structure or across an ensemble of structures from molecular dynamics trajectories.

## Data availability statement

All the basic data used to perform the simulations are freely available in Zenodo at https://doi.org/10.5281/zenodo.20506800.

## Supporting Information

Details of the comprehensive methodological details, as well as Figures S1–S5 and Table S1, S2 are provided in the Supporting Information. This material is available free of charge.

## Acknowledgement

The work conducted in Prof. Shengli Zhang’s group was funded by the National Natural Science Foundation of China (NSFC) under grant numbers T2425029, 12574231, 12304255. X Yu acknowledge support by Excellent Young Scientists Fund [Overseas] of China. The simulations were performed on the HPC platform of Xi’an Jiaotong University.

## REFERENCES

1. Mastroberardino L, Spindler B, Pfeiffer R, Skelly PJ, Loffing J, Shoemaker CB, et al. Amino-acid transport by heterodimers of 4F2hc/CD98 and members of a permease family. Nature. 1998;395: 288–291. doi:10.1038/26246

2. Fotiadis D, Kanai Y, Palacín M. The SLC3 and SLC7 families of amino acid tranporters. Mol Aspects Med. 2013;34: 139–158. doi:10.1016/j.mam.2012.10.007

3. Pfeiffer R, Spindler B, Loffing J, Skelly PJ, Shoemaker CB, Verrey F. Functional heterodimeric amino acid transporters lacking cysteine residues involved in disulfide bond. FEBS Lett. 1998;439: 157–162. doi:10.1016/s0014-5793(98)01359-3

4. Yan R, Zhao X, Lei J, Zhou Q. Structure of the human LAT1–4F2hc heteromeric amino acid transporter complex. Nature. 2019;568: 127–130. doi:10.1038/s41586-019-1011-z

5. Yan R, Li Y, Shi Y, Zhou J, Lei J, Huang J, et al. Cryo-EM structure of the human heteromeric amino acid transporter b0,+AT-rBAT. Sci Adv. 2020;6: eaay6379. doi:10.1126/sciadv.aay6379

6. Wu D, Grund TN, Welsch S, Mills DJ, Michel M, Safarian S, et al. Structural basis for amino acid exchange by a human heteromeric amino acid transporter. Proc Natl Acad Sci U S A. 2020;117: 21281–21287. doi:10.1073/pnas.2008111117

7. Sheng L, Luo Q, Chen L. Amino Acid Solute Carrier Transporters in Inflammation and Autoimmunity. Drug Metab Dispos. 2022;50: DMD-AR-2021-000705. doi:10.1124/dmd.121.000705

8. Errasti-Murugarren E, Palacín M. Heteromeric Amino Acid Transporters in Brain: from Physiology to Pathology. Neurochem Res. 2022;47: 23–36. doi:10.1007/s11064-021-03261-w

9. Yang H, Shi T, Dong J, Zhang T, Li Y, Guo Y, et al. Structural insights into the substrate transport mechanism of the amino acid transporter complex. J Biol Chem. 2025;301: 110569. doi:10.1016/j.jbc.2025.110569

10. Dai L, Zeng Q, Zhang T, Zhang Y, Shi Y, Li Y, et al. Structural basis for the substrate recognition and transport mechanism of the human y^+^ LAT1-4F2hc transporter complex. Sci Adv. 2025;11: eadq0558. doi:10.1126/sciadv.adq0558

11. Reig N, Chillarón J, Bartoccioni P, Fernández E, Bendahan A, Zorzano A, et al. The light subunit of system b(o,+) is fully functional in the absence of the heavy subunit. EMBO J. 2002;21: 4906–4914. doi:10.1093/emboj/cdf500

12. Bröer S, Palacín M. The role of amino acid transporters in inherited and acquired diseases. Biochem J. 2011;436: 193–211. doi:10.1042/BJ20101912

13. Brons A-K, Henthorn PS, Raj K, Fitzgerald CA, Liu J, Sewell AC, et al. SLC3A1 and SLC7A9 mutations in autosomal recessive or dominant canine cystinuria: a new classification system. J Vet Intern Med. 2013;27: 1400–1408. doi:10.1111/jvim.12176

14. Yan R, Li Y, Shi Y, Zhou J, Lei J, Huang J, et al. Cryo-EM structure of the human heteromeric amino acid transporter b0,+AT-rBAT. Sci Adv. 2020;6: eaay6379. doi:10.1126/sciadv.aay6379

15. Fernández E, Jiménez-Vidal M, Calvo M, Zorzano A, Tebar F, Palacín M, et al. The structural and functional units of heteromeric amino acid transporters. The heavy subunit rBAT dictates oligomerization of the heteromeric amino acid transporters. J Biol Chem. 2006;281: 26552–26561. doi:10.1074/jbc.M604049200

16. Lee Y, Wiriyasermkul P, Jin C, Quan L, Ohgaki R, Okuda S, et al. Cryo-EM structure of the human L-type amino acid transporter 1 in complex with glycoprotein CD98hc. Nat Struct Mol Biol. 2019;26: 510–517. doi:10.1038/s41594-019-0237-7

17. Chovancova E, Pavelka A, Benes P, Strnad O, Brezovsky J, Kozlikova B, et al. CAVER 3.0: A Tool for the Analysis of Transport Pathways in Dynamic Protein Structures. PLOS Computational Biology. 2012;8: e1002708. doi:10.1371/journal.pcbi.1002708

18. Pavelka A, Sebestova E, Kozlikova B, Brezovsky J, Sochor J, Damborsky J. CAVER: Algorithms for Analyzing Dynamics of Tunnels in Macromolecules. IEEE/ACM Trans Comput Biol Bioinform. 2016;13: 505–517. doi:10.1109/TCBB.2015.2459680

19. Bouysset C, Fiorucci S. ProLIF: a library to encode molecular interactions as fingerprints. J Cheminform. 2021;13: 72. doi:10.1186/s13321-021-00548-6

20. Girvan M, Newman MEJ. Community structure in social and biological networks. Proceedings of the National Academy of Sciences. 2002;99: 7821–7826. doi:10.1073/pnas.122653799

21. Csardi G, Nepusz T. The igraph software package for complex network research. Interjournal Complex Systems. 2006;2006. Available: http://www.semanticscholar.org/paper/1d2744b83519657f5f2610698a8ddd177ced4f5c

22. Sethi A, Eargle J, Black AA, Luthey-Schulten Z. Dynamical networks in tRNA:protein complexes. Proc Natl Acad Sci U S A. 2009;106: 6620–6625. doi:10.1073/pnas.0810961106

23. Gasper PM, Fuglestad B, Komives EA, Markwick PRL, McCammon JA. Allosteric networks in thrombin distinguish procoagulant vs. anticoagulant activities. Proc Natl Acad Sci U S A. 2012;109: 21216–21222. doi:10.1073/pnas.1218414109

24. Scarabelli G, Grant BJ. Mapping the Structural and Dynamical Features of Kinesin Motor Domains. PLOS Computational Biology. 2013;9: e1003329. doi:10.1371/journal.pcbi.1003329

25. Yao X-Q, Grant BJ. Domain-opening and dynamic coupling in the α-subunit of heterotrimeric G proteins. Biophys J. 2013;105: L08-10. doi:10.1016/j.bpj.2013.06.006

26. Van Wart AT, Durrant J, Votapka L, Amaro RE. Weighted Implementation of Suboptimal Paths (WISP): An Optimized Algorithm and Tool for Dynamical Network Analysis. J Chem Theory Comput. 2014;10: 511–517. doi:10.1021/ct4008603

27. Yao X-Q, Malik RU, Griggs NW, Skjærven L, Traynor JR, Sivaramakrishnan S, et al. Dynamic Coupling and Allosteric Networks in the α Subunit of Heterotrimeric G Proteins. J Biol Chem. 2016;291: 4742–4753. doi:10.1074/jbc.M115.702605

28. Reyes N, Ginter C, Boudker O. Transport mechanism of a bacterial homologue of glutamate transporters. Nature. 2009;462: 880–885. doi:10.1038/nature08616

29. Lee C, Kang HJ, von Ballmoos C, Newstead S, Uzdavinys P, Dotson DL, et al. A two-domain elevator mechanism for sodium/proton antiport. Nature. 2013;501: 573–577. doi:10.1038/nature12484

30. Yan N. Structural Biology of the Major Facilitator Superfamily Transporters. Annu Rev Biophys. 2015;44: 257–283. doi:10.1146/annurev-biophys-060414-033901

31. Jo S, Kim T, Im W. Automated Builder and Database of Protein/Membrane Complexes for Molecular Dynamics Simulations. PLOS ONE. 2007;2: e880. doi:10.1371/journal.pone.0000880

32. Jo S, Kim T, Iyer VG, Im W. CHARMM-GUI: A web-based graphical user interface for CHARMM. Journal of Computational Chemistry. 2008;29: 1859–1865. doi:10.1002/jcc.20945

33. Jo S, Lim JB, Klauda JB, Im W. CHARMM-GUI Membrane Builder for Mixed Bilayers and Its Application to Yeast Membranes. Biophysical Journal. 2009;97: 50–58. doi:10.1016/j.bpj.2009.04.013

34. Wu EL, Cheng X, Jo S, Rui H, Song KC, Dávila-Contreras EM, et al. CHARMM-GUI Membrane Builder toward realistic biological membrane simulations. Journal of Computational Chemistry. 2014;35: 1997–2004. doi:10.1002/jcc.23702

35. Lee J, Cheng X, Swails JM, Yeom MS, Eastman PK, Lemkul JA, et al. CHARMM-GUI Input Generator for NAMD, GROMACS, AMBER, OpenMM, and CHARMM/OpenMM Simulations Using the CHARMM36 Additive Force Field. J Chem Theory Comput. 2016;12: 405–413. doi:10.1021/acs.jctc.5b00935

36. Lee J, Patel DS, StÅhle J, Park S-J, Kern NR, Kim S, et al. CHARMM-GUI Membrane Builder for Complex Biological Membrane Simulations with Glycolipids and Lipoglycans. J Chem Theory Comput. 2019;15: 775–786. doi:10.1021/acs.jctc.8b01066

37. Lee J, Hitzenberger M, Rieger M, Kern NR, Zacharias M, Im W. CHARMM-GUI supports the Amber force fields. The Journal of Chemical Physics. 2020;153: 035103. doi:10.1063/5.0012280

38. Park S, Choi YK, Kim S, Lee J, Im W. CHARMM-GUI Membrane Builder for Lipid Nanoparticles with Ionizable Cationic Lipids and PEGylated Lipids. J Chem Inf Model. 2021;61: 5192–5202. doi:10.1021/acs.jcim.1c00770

39. Izadi S, Anandakrishnan R, Onufriev AV. Building Water Models: A Different Approach. J Phys Chem Lett. 2014;5: 3863–3871. doi:10.1021/jz501780a

40. Case DA, Aktulga HM, Belfon K, Cerutti DS, Cisneros GA, Cruzeiro VWD, et al. AmberTools. J Chem Inf Model. 2023;63: 6183–6191. doi:10.1021/acs.jcim.3c01153

41. Dickson CJ, Walker RC, Gould IR. Lipid21: Complex Lipid Membrane Simulations with AMBER. J Chem Theory Comput. 2022;18: 1726–1736. doi:10.1021/acs.jctc.1c01217

42. Tian C, Kasavajhala K, Belfon KAA, Raguette L, Huang H, Migues AN, et al. ff19SB: Amino-Acid-Specific Protein Backbone Parameters Trained against Quantum Mechanics Energy Surfaces in Solution. J Chem Theory Comput. 2020;16: 528–552. doi:10.1021/acs.jctc.9b00591

43. Götz AW, Williamson MJ, Xu D, Poole D, Le Grand S, Walker RC. Routine Microsecond Molecular Dynamics Simulations with AMBER on GPUs. 1. Generalized Born. J Chem Theory Comput. 2012;8: 1542–1555. doi:10.1021/ct200909j

44. Salomon-Ferrer R, Götz AW, Poole D, Le Grand S, Walker RC. Routine Microsecond Molecular Dynamics Simulations with AMBER on GPUs. 2. Explicit Solvent Particle Mesh Ewald. J Chem Theory Comput. 2013;9: 3878–3888. doi:10.1021/ct400314y

45. Le Grand S, Götz AW, Walker RC. SPFP: Speed without compromise—A mixed precision model for GPU accelerated molecular dynamics simulations. Computer Physics Communications. 2013;184: 374–380. doi:10.1016/j.cpc.2012.09.022

46. Roe DR, Cheatham TEI. PTRAJ and CPPTRAJ: Software for Processing and Analysis of Molecular Dynamics Trajectory Data. J Chem Theory Comput. 2013;9: 3084–3095. doi:10.1021/ct400341p

47. Jarzynski C. Nonequilibrium Equality for Free Energy Differences. Phys Rev Lett. 1997;78: 2690–2693. doi:10.1103/PhysRevLett.78.2690

48. Park S, Schulten K. Calculating potentials of mean force from steered molecular dynamics simulations. J Chem Phys. 2004;120: 5946–5961. doi:10.1063/1.1651473

49. Ozer G, Valeev EF, Quirk S, Hernandez R. Adaptive Steered Molecular Dynamics of the Long-Distance Unfolding of Neuropeptide Y. J Chem Theory Comput. 2010;6: 3026–3038. doi:10.1021/ct100320g

50. Ozer G, Quirk S, Hernandez R. Adaptive steered molecular dynamics: Validation of the selection criterion and benchmarking energetics in vacuum. J Chem Phys. 2012;2012. doi:10.1063/1.4725183

51. Ozer G, Quirk S, Hernandez R. Thermodynamics of Decaalanine Stretching in Water Obtained by Adaptive Steered Molecular Dynamics Simulations. J Chem Theory Comput. 2012;8: 4837–4844. doi:10.1021/ct300709u

52. Ozer G, Keyes T, Quirk S, Hernandez R. Multiple branched adaptive steered molecular dynamics. J Chem Phys. 2014;2014. doi:10.1063/1.4891807

53. Bureau HR, Jr DRM, Hershkovits E, Quirk S, Hernandez R. Constrained Unfolding of a Helical Peptide: Implicit versus Explicit Solvents. PLOS ONE. 2015;10: e0127034. doi:10.1371/journal.pone.0127034

54. Duan H, Hu K, Zheng D, Cheng Y, Zhang Z, Wang Y, et al. Recognition and release of uridine and hCNT3: From multivariate interactions to molecular design. International Journal of Biological Macromolecules. 2022;223: 1562–1577. doi:10.1016/j.ijbiomac.2022.11.145

55. Wang E, Sun H, Wang J, Wang Z, Liu H, Zhang JZH, et al. End-Point Binding Free Energy Calculation with MM/PBSA and MM/GBSA: Strategies and Applications in Drug Design. Chem Rev. 2019;119: 9478–9508. doi:10.1021/acs.chemrev.9b00055

56. Case DA, Cerutti DS, Cruzeiro VWD, Darden TA, Duke RE, Ghazimirsaeed M, et al. Recent Developments in Amber Biomolecular Simulations. J Chem Inf Model. 2025;65: 7835–7843. doi:10.1021/acs.jcim.5c01063

57. Shi S, Zhang S, Zhang Q. Probing Difference in Binding Modes of Inhibitors to MDMX by Molecular Dynamics Simulations and Different Free Energy Methods. PLoS One. 2015;10: e0141409. doi:10.1371/journal.pone.0141409

58. Wang L, Bao Q-C, Xu X-L, Jiang F, Gu K, Jiang Z-Y, et al. Discovery and identification of Cdc37-derived peptides targeting the Hsp90–Cdc37 protein–protein interaction. RSC Adv. 2015;5: 96138–96145. doi:10.1039/C5RA20408A

59. Yang Z, Wu F, Yuan X, Zhang L, Zhang S. Novel binding patterns between ganoderic acids and neuraminidase: Insights from docking, molecular dynamics and MM/PBSA studies. J Mol Graph Model. 2016;65: 27–34. doi:10.1016/j.jmgm.2016.02.006

60. Zang Y, Li X, Zhao Y, Wang H, Hao D, Zhang L, et al. Molecular insights into the binding variance of the SARS-CoV-2 spike with human, cat and dog ACE2 proteins. Phys Chem Chem Phys. 2021;23: 13752–13759. doi:10.1039/d1cp01611c

61. Galindo-Murillo R, Roe DR, Cheatham TE. Convergence and reproducibility in molecular dynamics simulations of the DNA duplex d(GCACGAACGAACGAACGC). Biochimica et Biophysica Acta (BBA) - General Subjects. 2015;1850: 1041–1058. doi:10.1016/j.bbagen.2014.09.007

62. Abraham MJ, Murtola T, Schulz R, Páll S, Smith JC, Hess B, et al. GROMACS: High performance molecular simulations through multi-level parallelism from laptops to supercomputers. SoftwareX. 2015;1–2: 19–25. doi:10.1016/j.softx.2015.06.001

63. Bouysset C, Fiorucci S. ProLIF: a library to encode molecular interactions as fingerprints. Journal of Cheminformatics. 2021;13: 72. doi:10.1186/s13321-021-00548-6

